# Sporulation, bacterial cell envelopes, and the origin of life

**DOI:** 10.1101/041624

**Authors:** Elitza I. Tocheva, Davi R. Ortega, Grant J. Jensen

**Author notes:** For correspondence. Grant J. Jensen.

## Abstract

Four recent papers from our group exploiting the power of electron cryotomography to produce 3-D reconstructions of intact cells in a near-native state have led to the proposal that an ancient sporulation-like event gave rise to the second membrane in diderm bacteria. Here we review the images of sporulating monoderm and diderm cells which show how sporulation leads to diderm cells. We also review the images of Gram-negative and Gram-positive cell walls that show they are more closely related than previously thought, and explain how this provides critical support for the hypothesis. Mapping the distribution of cell envelope architectures onto the most recent phylogenetic tree of life then leads to the conclusion that the diderm cell plan, and therefore the sporulation-like event that gave rise to it, must be very ancient. One explanation for the biogeologic record is that during the cataclysmic transitions of early Earth, cellular evolution may have gone through a bottleneck where only spores survived (LUCA was a spore).

## Introduction

Historically, bacteria were first classified by their ability to retain the Gram stain^1^. With the development of electron microscopy, the actual structure of the bacterial cell envelope became apparent. Gram-positive bacteria were seen to be monoderms (possessing a single membrane) with a relatively thick peptidoglycan (PG) layer. Gram-negative bacteria were seen to be diderms (possessing two membranes) with a thin PG layer between the two membranes. Typical Gram-positive and Gram-negative cell envelope architectures are exhibited by the phyla *Firmicutes* and *Proteobacteria* and represented by the model organisms *Bacillus subtilis* and *Escherichia coli*, respectively^2^.

The cytoplasmic membrane of all cells (the innermost in the case of diderm bacteria) is loaded with α-helical proteins and sustains a chemical (usually proton) gradient used to make ATP. The structure and function of the outer membrane (OM) in diderm bacteria is very different. While the inner membrane (IM) is a symmetrical lipid bilayer, the OM of typical Gram-negative bacteria is asymmetric, with the outer leaflet composed primarily of lipopolysaccharide (LPS). OMs are also rich in outer membrane proteins (OMPs, mostly β-barrels) that allow free diffusion of small molecules in and out of the periplasm^3^. Interesting exceptions to the typical Gram-positive and-negative envelope architectures exist. For example, *Mycoplasma* (phylum *Tenericutes*) lack PG and *Mycobacteria*, a genus of the phylum *Actinobacteria*, have an unusual OM rich in mycolic acids.

## Outer membrane biogenesis through sporulation

Various theories about the origin of life on Earth exist. Some imagine larger and larger molecules replicating in rich organic pools, with lipid vesicles washing out of cavities in rocks to enclose primordial cells^4^. Assuming that the earliest primordial cells were monoderm ^2, 5-7^, the acquisition of an OM must have been a major evolutionary step. Prior to a paper we published in 2011^8^, the most prominent hypothesis for how OMs evolved was based on distributions of protein families and proposed that a symbiosis between an ancient actinobacterium and an ancient clostridium produced the last common ancestor of all diderms^9^. Our work suggested a fundamentally different mechanism based on endospore formation, a common process executed by many *Firmicute* species. Endospore formation begins with genome segregation and asymmetric cell division. Next, in a process similar to phagocytosis in eukaryotic cells, the bigger compartment (mother cell) engulfs the smaller compartment (future spore). Multiple protective layers are formed around the immature spore, and then lysis of the mother cell releases a mature spore into the environment. Notably, spores have a complete copy of the species’ genomic DNA. Later, germination returns the dormant spore to a vegetative state^10^.

While the well-studied model endospore-forming species (classes *Bacilli* and *Clostridia*) are all monoderms, we imaged *Acetonema longum*, a member of a lesser-known family of *Clostridia* named the *Veillonellaceae* that forms endospores but is diderm^8^. Biochemical characterization of the outer membrane revealed that it contained LPS just like model diderms. Homologies between many *A. longum* OMPs and their counterparts in *Proteobacteria* and other diderm species suggested these proteins share an ancient common heritage (rather than being the result of convergent evolution)^8^. Moreover, phylogenetic analysis of several Omp85/Omp87 clearly revealed the close relationship between mitochondria and α-proteobacteria, for example, but no special relationship between *A. longum* and any other diderm phylum, arguing against recent horizontal gene transfer^8^. *A. longum* and other members of *Veillonellaceae* are therefore candidate missing links between monoderm and diderm bacteria since they possess characteristics of both: an OM and the ability to form endospores.

Mechanistic clues about how endospore formation may have given rise to bacterial OMs came from comparing images of sporulating *B. subtilis* (monoderm) and *A. longum* (diderm) cells (Figure 1) ^8,11^. Each were imaged with electron cryotomography (ECT), which provides 3-dimensional reconstructions of intact cells to “macromolecular” (3-4 nm) resolution ^12^. Images of vegetative, sporulating and germinating cells revealed that both monoderm (*B. subtilis*, Figure 1a-f) and diderm (*A. longum*, Figure 1a’-f’) bacteria produced spores that were surrounded by two membranes. Furthermore, in both cases the two membranes originated from the inner/cytoplasmic membrane of the mother cell. Some time between mid to late spore development and germination, *B. subtilis* loses its outer spore membrane to become a monoderm, “Gram-positive” vegetative cell, whereas *A. longum* retains both spore membranes and, amazingly, the outer spore membrane emerges as an OM. Therefore, endospore formation offers a novel hypothesis for how the bacterial OM could have evolved: a primordial monoderm cell may have first developed the ability to form endospores, and then this process could have given rise to diderm vegetative cells (Figure 2).

**Figure 1.**
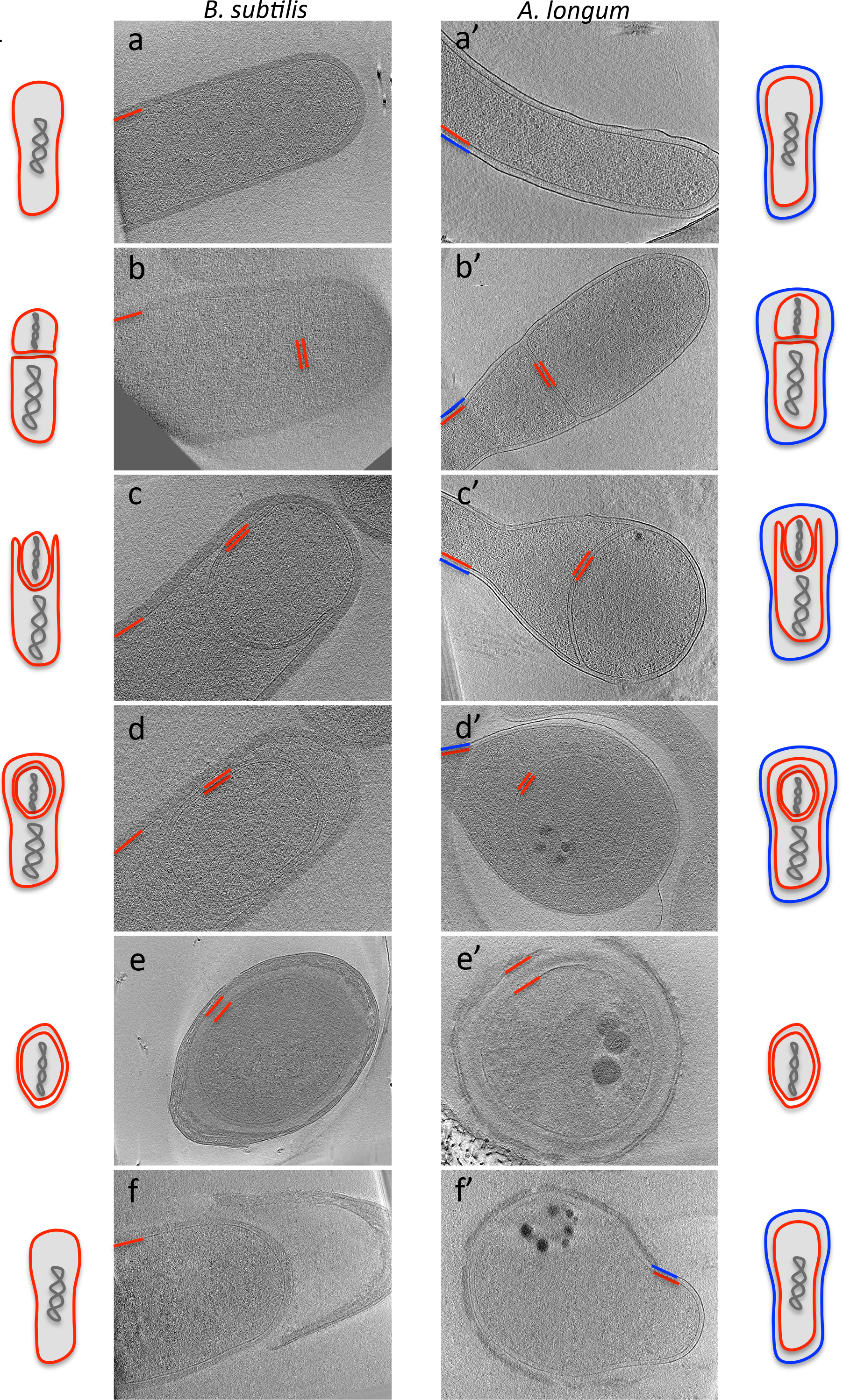
Sporulation in Gram-positive and Gram-negative bacteria. Overview of bacterial sporulation in *B. subtilis* (a-f) and *A. longum* (a’-f’) by electron cryotomography. Each panel represents a tomographic slice through a bacterial cell at a different stage of sporulation. The inner and cytoplasmic membranes are depicted in red and the OM of *A. longum* is depicted in blue. Schematic representations of all sporulation stages are shown next to the tomographic slices.

**Figure 2.**
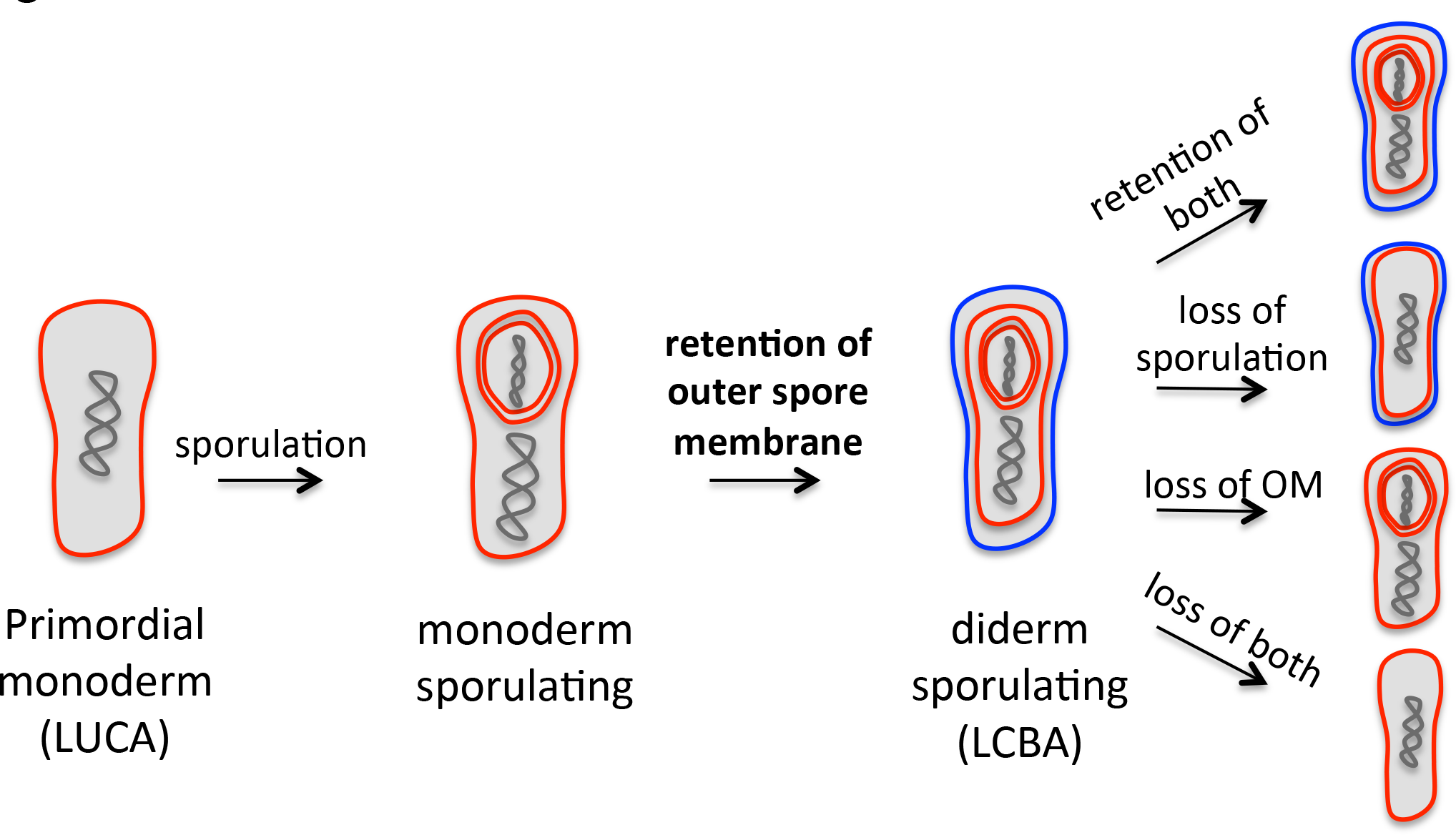
Proposed mechanism of how sporulation gave rise to OM in bacteria. At some early point in evolution, the cell division and nutrient-uptake processes of a primordial cell were combined into a sporulation-like process. Retention of the second spore membrane led to diderm sporulating species. Losses of the OM, the ability to sporulate or both in various lineages can explain the distribution of cell envelope architectures seen in modern bacteria.

## Peptidoglycan architecture and remodeling during sporulation

Further support for this hypothesis came from our imaging studies of the cell wall in both Gram-positive and Gram-negative species. Typical Gram-negative PG is a single-layer polymer composed of long glycan strands formed by repeating units of *N-*acetyl glucosamine:*N*-acetyl muramic acid cross-linked with peptide bonds^13^. PG is synthesized by transglycosylases and transpeptidases and other proteins, influenced by cytoskeletal filaments^14^. The architecture of thin, Gram-negative PG was initially unclear. Although various indirect lines of evidence favored models in which the glycan strands run parallel to the cell surface (“layered” model), alternative models such as a perpendicular, “scaffold” model, had been proposed^15-17^. Direct visualization of the architecture of PG in *E. coli* and *Caulobacter crescentus* by ECT revealed that the glycan strands of Gram-negative PG run along the cell surface perpendicular to the long axis of the cell (the “layered” model, which we prefer to call “circumferential”^18^).

Gram-positive PG is much thicker (~40 nm), but is known to be closely related to Gram-negative PG for two reasons. First, the basic chemical structure of PG is very similar among Gram-positive and Gram-negative bacteria – notable differences are mostly related to modifications in the peptide composition, the degree of peptide crosslinks, and the length of the peptidoglycan chains^19^. In fact, *Proteobacteria* and *Firmicutes*, in particular, even share the same chemotype of PG (A1γ), with a *meso*-A_2_pm residue at position 3 of the peptide and a direct crosslink to a D-Ala at position 4 of the neighboring peptide^20,21^. Second, numerous biochemical and genetic studies have shown that the enzymes responsible for synthesizing PG in Gram-negative and Gram-positive species are homologous^22-24^, so the cell walls they build must also be similar.

A variety of models for the architecture of Gram-positive PG have been proposed including circumferential and scaffold but also an additional, “coiled cable” model^16,25-28^. Unfortunately, unlike Gram-negative PG, direct visualization of the architecture of Gram-positive PG by ECT was not possible due to its thickness and rigidity^29^. ECT imaging of sheared purified *B. subtilis* sacculi and coarse grain molecular dynamics simulations were able however to refute the coiled cable and scaffold models and uniquely support the “circumferential” model^29^. Thus the basic architecture of both Gram-negative and-positive cell walls are the same.

Interestingly, our studies of sporulating *B. subtilis* (Gram-positive, thick PG) and *A. longum* (Gram-negative, thin PG) cells confirmed that Gram-negative and Gram-positive PG have the same basic architecture by revealing that they can be interconverted^11^. At the onset of sporulation in *B. subtilis*, thick PG (~40 nm) is present between the two septal membranes (Figure 3, top). This thick PG is then remodeled into a thin, Gram-negative-like PG prior to engulfment. The thin layer is extended by the synthesis of new PG at the leading edges of engulfing membranes as they progress around the immature spore^30^. At the end of engulfment, a thin layer of PG is found between the two spore membranes and likely acts as a foundation for the synthesis of the spore cortex (thick protective layers of glycan strands cross-linked by peptide bonds). Similar transitions of PG thickness were observed in *A. longum* (Figure 3, bottom). Thus, our tomograms showed that both *A. longum* and *B. subtilis* transform a thick PG layer into a thin, Gram-negative-like PG layer that eventually surrounds the immature spore during engulfment, and then expand this thin layer into a thick cortex during spore maturation. In other words, our ECT data show that both Gram-negative and Gram-positive bacteria can synthesize both thin and thick PG and can gradually remodel one into the other, strongly confirming the concept that the cell walls have the same basic architecture (circumferential), and must differ mainly in just the number of layers present^11^. The existence of a variety of PG thicknesses in other species such as the presence of medium-thick PG in cyanobacteria further support this conclusion^31^.

**Figure 3.**
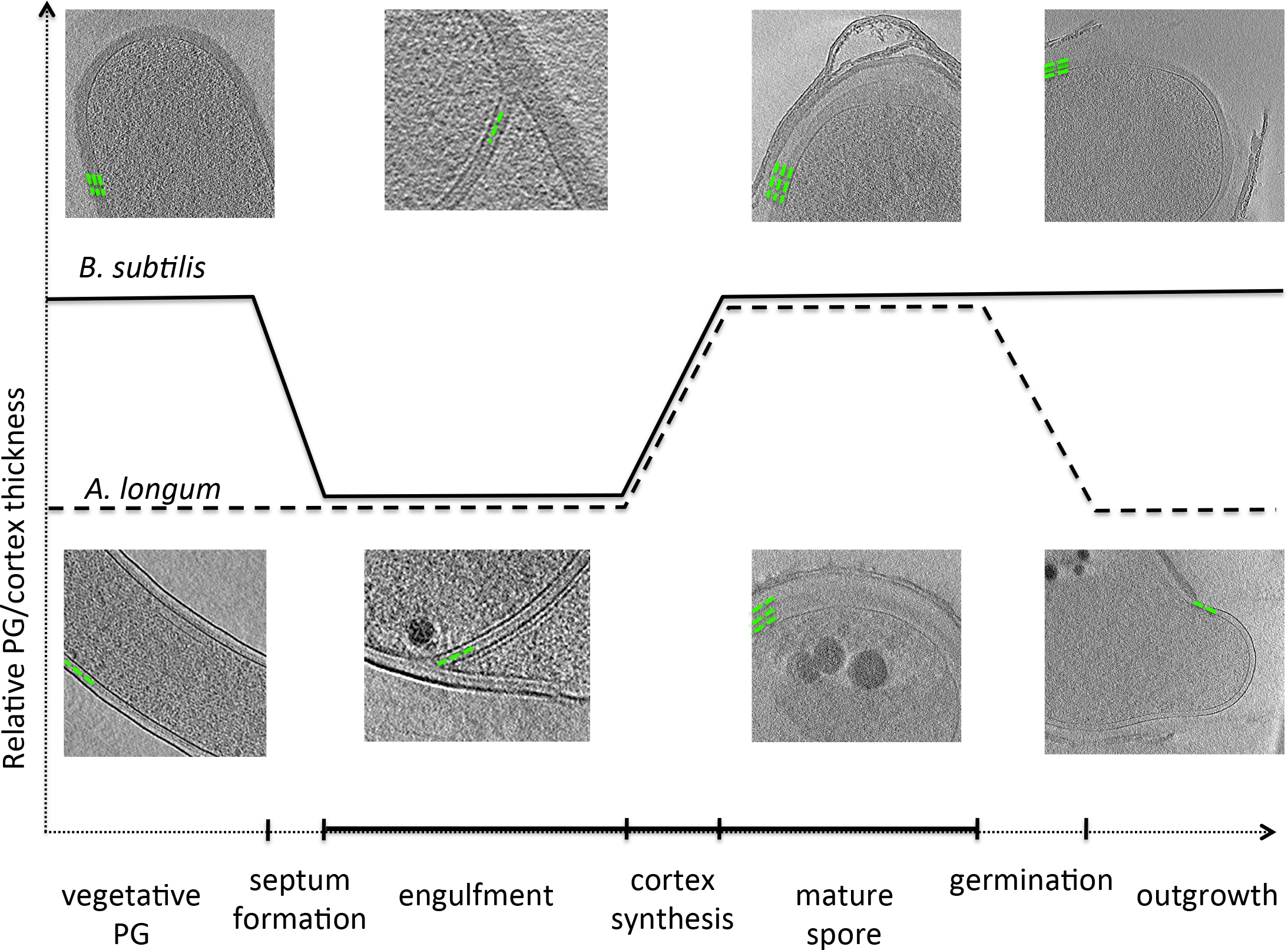
Peptidoglycan remodeling during sporulation. Peptidoglycan and cortex remodeling during sporulation in *B. subtilis* and *A. longum* shows that both Gram-positive and Gram-negative cells can synthesize thick and thin PG and remodel one into the other, supporting the notion that all bacterial cell walls have the same basic architecture inherited from a common ancestor. Adapted from Tocheva et al Mol. Microbiol. 2013 (with permission).

Because Gram-negative and-positive cell walls have the same architecture and are interconvertible, we should not think of these two major divisions of bacteria as completely separate branches of the bacterial tree, but instead as potentially closely related or even phylogenetically intermixed (as they are in the *Firmicute* phylum^32,33^). This supports the hypothesis that the sporulation process in a primordial cell may have led to both the Gram-positive and-negative cell plans.

## Evolutionary implications

As the number of sequenced genomes has increased, more and more sophisticated phylogenetic analyses of Bacteria have been possible^3,34-36^. While this has made the relationships between phyla increasingly clear, unfortunately there is no agreement on how to root the tree of life, so it remains a mystery which modern species most closely resembles the last universal common ancestor (LUCA)^37^. In order to explore the evolutionary implications of our hypothesis, here we consider three different roots for the tree of life: i) between Archaea and Bacteria^38,39^ (Figure 4A) (even though historically most favored, it is not supported by the current state of knowledge^40^); ii) at the phylum Chloroflexi^41,42^ (Figure 4B); and iii) at the phylum Firmicutes^3,43^ (Figure 4C) (the last two roots are suggested by current methodologies in systematics). In each case we use the most recent and comprehensive published tree of life^34^ to assert relationships between phyla, simply rooting it in different places, and we map the gains and losses of the ability to sporulate and the presence of an outer membrane that minimize the number of grand evolutionary events required (Figure 4).

**Figure 4.**
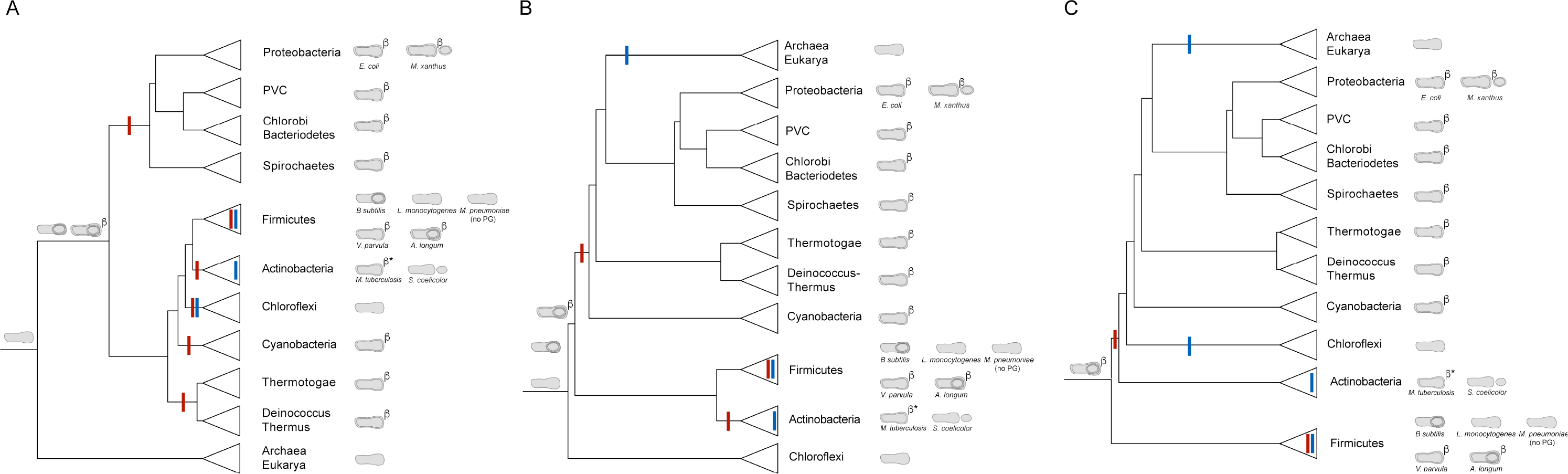
Rooted phylogenetic trees representing relationships between bacterial phyla. Three different schematic reproductions of the most recent phylogenetic tree of the major bacterial phyla based on 42 gene markers conserved throughout Bacteria and Archaea rooted in three different ways, following leading proposals in the field. Cell envelope structures exhibited by members of these phyla are shown on the right. Reported abilities to sporulate are represented by small circles within the cells (for endospores) or next to them (for exospores). The presence of conserved Omp85/Omp87 is noted by the “β” symbol and the presence of other conserved OMPs is noted by “β*”. The deduced basic cell plans of ancient ancestors are depicted on early branches.

We begin with the first scenario (that the root of the tree of life lies between the three major kingdoms), as suggested by Gogarten and Woese^37,38^. Mapping the basic cell envelope structures of different species to the right of the corresponding tree (Figure 4A) presents an interesting surprise: the monoderm phyla (*Firmicutes*, *Tenericutes*, *Actinobacteria*, and *Chloroflexi*) are surrounded by diderms. In addition, the *Firmicutes* and *Actinobacteria* phyla comprise both monoderm and diderm species. Unless the diderm cell plan evolved multiple times independently, or was horizontally transferred (both highly unlikely, since the OM structure depends on hundreds of genes and OM proteins are homologous among all diderm phyla), this suggests that the last bacterial common ancestor (LBCA) was a diderm. The diversity of known phyla could then be explained by losses of either the OM and/or sporulation properties or both: diderm phyla such as *Deinococcus-Thermus*, *Thermotoga*, *Cyanobacteria*, *Spirochaetes*, *Chlorobi-Bacteriodetes*, and *Planctomycetes-Verrucomicrobia-Chlamydiae* (PVC superphylum) lost their ability to sporulate but retained their OM (explaining why they all share common β-barrel OMPs ^8,44^). *Proteobacteria* retained the OM and lost the ability to sporulate. Within *Firmicutes*, most *Bacilli* and *Clostridia* (like *B. subtilis* and *Clostridia difficile*) retained their ability to form endospores but discarded their OM in the vegetative state, perhaps for reasons of increased efficiency. Other *Clostridia* like *A. longum* retained both properties while still others such as *Veillonella parvula* lost the ability to sporulate but retained an OM. *Chloroflexi* and some *Firmicutes* (such as *Listeria monocytogenes*) lost both the ability to sporulate and their OM, as did the *Tenericutes* (*Mycoplasma* spp.), which further discarded their PG.

The *Actinobacteria* phylum presents a particularly interesting case. Some *Actinobacteria* like *Streptomyces coelicolor* are monoderm. Others like *Mycobacteria* are diderm, but have a unique outer membrane linked to a thin PG layer via an arabinogalactan network^45^. The lipid composition of the outer membrane is rich in mycolic acid, a notable difference from the OM of *Proteobacteria*. Even though different lipids comprise the outer membrane in *Mycobacteria*, bioinformatics and experimental approaches have identified numerous OMPs with β-barrel structure homologous to the OMPs in *Proteobacteria* ^46-48^. Mycobacterial OMPs have even been shown to have the same signature sequences as their Proteobacterial homologs^49^. The simplest explanation is that *Actinobacteria* descend from an ancestor with an OM and established OMPs but the OM lipids in the Mycobacterial branch were exchanged for mycolic acids.

The second evolutionary scenario we will consider arises from the proposals of Cavalier-Smith and Valas and Bourne^40,41^ that modern *Chloroflexi* are most closely related to the root of the tree of life (Figure 4B). In this scenario, the *Chloroflexi* may have never been diderm, but for similar reasoning as above (it is unlikely that the diderm cell plan evolved multiple times or was laterally transferred), a very early sporulation process must have led to a diderm that is the last common ancestor of all other cells (in bioinformatics parlance, we argue this because all the diderms are not monophyletic). A third scenario (Figure 4C) is that modern *Firmicutes* are closest to the root (Lake and Bork^3,43^). Again this would imply that one monoderm Firmicute branch may never have been diderm, but assuming the diderm cell plan evolved only once, a very early sporulation event led to the diderm common ancestor of all other cells, and the diversity seen today in the rest of the tree is the result of losses.

## Conclusions

In summary, we have presented the hypothesis that the diderm cell plan arose from a sporulation-like event. We have reviewed the images of sporulating monoderm and diderm species and Gram-negative and Gram-positive cell walls that support the hypothesis. We have considered the implications of this hypothesis in light of the increasingly well-characterized phylogenetic Tree of Life. While the root of the tree is still unknown, there are two conclusions that can be made regardless of which root is correct: (1) most if not all monoderms have not always been monoderm, but rather are the result of a diderm losing its OM; (2) sporulation and the diderm plan are extremely ancient, preceding all or at most one phylum-level branch point.

Early Earth history remains unclear, but we argue that one of the very first cells was a endospore-former. Interestingly, spores have long been recognized as exquisitely robust life forms and are known to be able to withstand all kinds of environmental insults such as temperature and pH extremes and dehydration^50^. One possible explanation of the biogeologic record is that during the cataclysmic early conditions on Earth, cellular evolution went through a bottleneck in which only robust spores survived, so LUCA was a spore. An alternative explanation is panspermia^51^: life *arrived* on Earth as a spore, for instance in one of the frozen comets that some theorize created the oceans. As the Earth became more clement, for efficiency major branches of the bacterial tree lost the ability to sporulate, resulting in large numbers of diderm non-sporulating phyla today. Other explanations are also possible.

In any case, our analysis at least challenges the notion that complexity gradually increases through evolutionary time: instead it is a case of high complexity (a diderm endospore-forming cell) existing very early, followed by billions of years of losses with comparatively minor modification like the exchange of mycolic acids for LPS in the OM of *Mycobacteria*. This agrees with other recent evidence that complexity increased rapidly in early life on Earth followed by a long period of simplification^52^. We acknowledge, however, that it will be difficult if not impossible to test these speculations, and that some investigators believe that genes have been swapped so frequently and in such numbers across so many species that trees of life don’t even make sense^53^. Nevertheless we believe our idea that the OM arose through a sporulation event is intriguing, and that if it did, parsimony argues it must have been a very ancient, perhaps even initiatory, event.

## Acknowledgements

This project/ publication was made possible through the support of a grant from the John Templeton Foundation. The opinions expressed in this publication are those of the authors and do not necessarily reflect the views of the John Templeton Foundation.

## References

1 Gram, C. Ueber die isolirte Firbung der Schizomyceten iu Schnitt-und Trockenpriparate. Fortschitte der Medicin 2, 185–189 (1884).

2 Gupta, R. S. Protein phylogenies and signature sequences: A reappraisal of evolutionary relationships among archaebacteria, eubacteria, and eukaryotes. Microbiology and molecular biology reviews: MMBR 62, 1435–1491 (1998).

3 Ciccarelli, F. D. et al. Toward automatic reconstruction of a highly resolved tree of life. Science 311, 1283–1287, doi:10.1126/science.1123061 (2006).

4 Martin, W., Baross, J., Kelley, D. & Russell, M. J. Hydrothermal vents and the origin of life. Nature reviews. Microbiology 6, 805–814, doi:10.1038/nrmicro1991 (2008).

5 Gupta, R. S. & Golding, G. B. Evolution of HSP70 gene and its implications regarding relationships between archaebacteria, eubacteria, and eukaryotes. Journal of molecular evolution 37, 573–582 (1993).

6 Koch, A. L. Were Gram-positive rods the first bacteria? Trends Microbiol 11, 166–170 (2003).

7 Lake, J. A., Herbold, C. W., Rivera, M. C., Servin, J. A. & Skophammer, R. G. Rooting the tree of life using nonubiquitous genes. Mol Biol Evol 24, 130–136, doi:10.1093/molbev/msl140 (2007).

8 Tocheva, E. I. et al. Peptidoglycan remodeling and conversion of an inner membrane into an outer membrane during sporulation. Cell 146, 799–812, doi:S0092-8674(11)00827-0 [pii]10.1016/j.cell.2011.07.029 (2011).

9 Lake, J. A. Evidence for an early prokaryotic endosymbiosis. Nature 460, 967–971, doi:nature08183 [pii]10.1038/nature08183 (2009).

10 Kay, D. & Warren, S. C. Sporulation in Bacillus subtilis. Morphological changes. Biochem J 109, 819–824 (1968).

11 Tocheva, E. I. et al. Peptidoglycan transformations during Bacillus subtilis sporulation. Mol Microbiol 88, 673–686, doi:10.1111/mmi.12201 (2013).

12 Gan, L. & Jensen, G. J. Electron tomography of cells. Quarterly reviews of biophysics 45, 27–56, doi:10.1017/S0033583511000102 (2012).

13 Typas, A., Banzhaf, M., Gross, C. A. & Vollmer, W. From the regulation of peptidoglycan synthesis to bacterial growth and morphology. Nature reviews. Microbiology 10, 123–136, doi:nrmicro2677 [pii]10.1038/nrmicro2677 (2011).

14 Vollmer, W. & Bertsche, U. Murein (peptidoglycan) structure, architecture and biosynthesis in Escherichia coli. Biochim Biophys Acta 1778, 1714–1734, doi:S0005-2736(07)00221-0 [pii]10.1016/j.bbamem.2007.06.007 (2008).

15 Verwer, R.W., Nanninga, N., Keck, W. & Schwarz, U. Arrangement of glycan chains in the sacculus of Escherichia coli. J Bacteriology 136, 723–729 (1978).

16 Dmitriev, B. A. et al. Tertiary structure of bacterial murein: the scaffold model. J Bacteriol 185, 3458–3468 (2003).

17 Vollmer, W. & Holtje, J. V. The architecture of the murein (peptidoglycan) in gram-negative bacteria: vertical scaffold or horizontal layer(s)? J Bacteriol 186, 5978–5987, doi:10.1128/JB.186.18.5978-5987.2004.186/18/5978 [pii] (2004).

18 Gan, L., Chen, S. & Jensen, G. J. Molecular organization of Gram-negative peptidoglycan. Proc Natl Acad Sci USA 105, 18953–18957,doi:10.1073/pnas.0808035105 (2008).

19 Vollmer, W., Blanot, D. & de Pedro, M. A. Peptidoglycan structure and architecture. FEMS Microbiol Rev 32, 149–167, doi:FMR094 [pii]10.1111/j.1574-6976.2007.00094.x (2008).

20 Schleifer, K. H. & Kandler, O. Peptidoglycan types of bacterial cell walls and their taxonomic implications. Bacteriol Rev 36, 407–477 (1972).

21 Vollmer, W. Bacterial outer membrane evolution via sporulation? Nature chemical biology 8, 14–18, doi:nchembio.748 [pii]10.1038/nchembio.748 (2011).

22 McPherson, D.C., Driks, A. & Popham, D. L. Two class A high-molecular-weight penicillin-binding proteins of Bacillus subtilis play redundant roles in sporulation. J Bacteriol 183, 6046–6053, doi:10.1128/JB.183.20.6046-6053.2001 (2001).

23 Sauvage, E., Kerff, F., Terrak, M., Ayala, J. A. & Charlier, P. The penicillin-binding proteins: structure and role in peptidoglycan biosynthesis. FEMS Microbiol Rev 32, 234–258, doi:FMR105 [pii]10.1111/j.1574-6976.2008.00105.x (2008).

24 Foster, S. & Popham, D. in Bacillus subtilis and its close relatives: from genes to cells (eds A. L. Sonenshein, J. A. Hoch, & R. Losick) 21–41 (American Society for Microbiology, 2002).

25 Verwer, R. W. & Nanninga, N. Electron microscopy of isolated cell walls of Bacillus subtilis var. niger. Arch Microbiol 109, 195–197 (1976).

26 Dominguez-Escobar, J. et al. Processive movement of MreB-associated cell wall biosynthetic complexes in bacteria. Science 333, 225–228, doi:10.1126/science.1203466 (2011).

27 Garner, E. C. et al. Coupled, circumferential motions of the cell wall synthesis machinery and MreB filaments in B. subtilis. Science 333, 222–225, doi:10.1126/science.1203285 (2011).

28 Hayhurst, E.J., Kailas, L., Hobbs, J. K. & Foster, S. J. Cell wall peptidoglycan architecture in Bacillus subtilis. Proceedings of the National Academy of Sciences of the United States of America 105, 14603–14608, doi:0804138105 [pii]10.1073/pnas.0804138105 (2008).

29 Beeby, M., Gumbart, J. C., Roux, B. & Jensen, G. J. Architecture and assembly of the Gram-positive cell wall. Mol Microbiol 88, 664–672, doi:10.1111/mmi.12203 (2013).

30 Meyer, P., Gutierrez, J., Pogliano, K. & Dworkin, J. Cell wall synthesis is necessary for membrane dynamics during sporulation of Bacillus subtilis. Mol Microbiol 76, 956–970 (2010).

31 Hoiczyk, E. & Hansel, A. Cyanobacterial cell walls: news from an unusual prokaryotic envelope. J Bacteriol 182, 1191–1199 (2000).

32 Vesth, T. et al. Veillonella, Firmicutes: Microbes disguised as Gram negatives. Standards in genomic sciences 9, 431–448, doi:10.4056/sigs.2981345 (2013).

33 Yutin, N. & Galperin, M. Y. A genomic update on clostridial phylogeny: Gram-negative spore formers and other misplaced clostridia. Environ Microbiol 15, 2631–2641, doi:10.1111/1462-2920.12173 (2013).

34 Raymann, K., Brochier-Armanet, C. & Gribaldo, S. The two-domain tree of life is linked to a new root for the Archaea. Proceedings of the National Academy of Sciences of the United States of America, doi:10.1073/pnas.1420858112 (2015).

35 Wu, D. et al. A phylogeny-driven genomic encyclopaedia of Bacteria and Archaea. Nature 462, 1056–1060, doi:10.1038/nature08656 (2009).

36 Puigbo, P., Wolf, Y. I. & Koonin, E. V. Seeing the Tree of Life behind the phylogenetic forest. BMC biology 11, 46, doi:10.1186/1741-7007-11-46 (2013).

37 Lake, J.A., Skophammer, R. G., Herbold, C. W. & Servin, J. A. Genome beginnings: rooting the tree of life. Philosophical transactions of the Royal Society of London. Series B, Biological sciences 364, 2177–2185, doi:364/1527/2177 [pii]10.1098/rstb.2009.0035 (2009).

38 Woese, C. R., Kandler, O. & Wheelis, M. L. Towards a natural system of organisms: proposal for the domains Archaea, Bacteria, and Eucarya. Proceedings of the National Academy of Sciences of the United States of America 87, 4576–4579 (1990).

39 Gogarten, J. P. et al. Evolution of the vacuolar H+-ATPase: implications for the origin of eukaryotes. Proceedings of the National Academy of Sciences of the United States of America 86, 6661–6665 (1989).

40 Gouy, R., Baurain, D. & Philippe, H. Rooting the tree of life: the phylogenetic jury is still out. Philosophical transactions of the Royal Society of London. Series B, Biological sciences 370, 20140329, doi:10.1098/rstb.2014.0329 (2015).

41 Cavalier-Smith, T. Rooting the tree of life by transition analyses. Biology direct 1, 19, doi:10.1186/1745-6150-1-19 (2006).

42 Valas, R. E. & Bourne, P. E. Structural analysis of polarizing indels: an emerging consensus on the root of the tree of life. Biology direct 4, 30, doi:10.1186/1745-6150-4-30 (2009).

43 Lake, J. A. & Sinsheimer, J. S. The deep roots of the rings of life. Genome biology and evolution 5, 2440–2448, doi:10.1093/gbe/evt194 (2013).

44 Sutcliffe, I. C. A phylum level perspective on bacterial cell envelope architecture. Trends Microbiol 18, 464–470, doi:10.1016/j.tim.2010.06.005 (2010).

45 Nigou, J., Gilleron, M., Brando, T. & Puzo, G. Structural analysis of mycobacterial lipoglycans. Appl Biochem Biotechnol 118, 253–267 (2004).

46 Hillmann, D., Eschenbacher, I., Thiel, A. & Niederweis, M. Expression of the major porin gene mspA is regulated in Mycobacterium smegmatis. J Bacteriol 189, 958–967, doi:10.1128/JB.01474-06 (2007).

47 Song, H., Sandie, R., Wang, Y., Andrade-Navarro, M. A. & Niederweis, M. Identification of outer membrane proteins of Mycobacterium tuberculosis. Tuberculosis 88, 526–544, doi:10.1016/j.tube.2008.02.004 (2008).

48 Remmert, M., Biegert, A., Linke, D., Lupas, A. N. & Soding, J. Evolution of outer membrane beta-barrels from an ancestral beta beta hairpin. Mol Biol Evol 27, 1348–1358, doi:10.1093/molbev/msq017 (2010).

49 Nikaido, H. Molecular basis of bacterial outer membrane permeability revisited. Microbiology and molecular biology reviews: MMBR 67, 593–656 (2003).

50 Setlow, P. I will survive: DNA protection in bacterial spores. Trends Microbiol 15, 172–180, doi:S0966-842X(07)00026-1 [pii]10.1016/j.tim.2007.02.004 (2007).

51 Nicholson, W. L. Ancient micronauts: interplanetary transport of microbes by cosmic impacts. Trends Microbiol 17, 243–250, doi:10.1016/j.tim.2009.03.004 (2009).

52 Wolf, Y. I. & Koonin, E. V. Genome reduction as the dominant mode of evolution. Bioessays 35, 829–837, doi:10.1002/bies.201300037 (2013).

53 Doolittle, W. F. Phylogenetic classification and the universal tree. Science 284, 2124–2129 (1999).

